# Mutations in *PIKFYVE* cause autosomal dominant congenital cataract

**DOI:** 10.1101/2021.06.25.449865

**Authors:** Shaoyi Mei, Yi Wu, Yan Wang, Yubo Cui, Miao Zhang, Tong Zhang, Xiaosheng Huang, Sejie Yu, Tao Yu, Jun Zhao

## Abstract

Congenital cataract, an ocular disease predominantly occurring within the first decade of life, is one of the leading causes of blindness in children. Through whole exome sequencing of a Chinese family with congenital cataract, we identified a disease-causing mutation (p.G1943E) in *PIKFYVE*, which affecting the PIP kinase domain of the PIKfyve protein. We demonstrated that heterozygous/homozygous disruption of PIKfyve kinase domain, instead of overexpression of *PIKFYVE^G1943E^* in zebrafish mimicked the cataract defect in human patients, suggesting that haploinsufficiency, rather than dominant-negative inhibition of PIKfyve activity caused the disease. Phenotypical analysis of *pikfyve* zebrafish mutants revealed that loss of Pikfyve caused aberrant vacuolation (accumulation of Rab7^+^Lc3^+^ amphisomes) in lens cells, which was significantly alleviated by treatment with the V-ATPase inhibitor bafilomycin A1 (Baf-A1). Collectively, we identified *PIKFYVE* as a novel causative gene for congenital cataract and demonstrated the potential application of Baf-A1 in treatment of congenital cataract.

## Introduction

Congenital cataract is partial or complete opacification of the lens that occurs at birth or during the first decade of life. It is a common ocular abnormality that causes visual impairment and blindness during infancy (*Khokhar et al., 2017*), accounting for 5%-20% blindness in children worldwide (*Sheeladevi, Lawrenson, Fielder, & Suttle, 2016*). According to the location and shape of the lens opacities, congenital cataract can be divided into seven clinical types: nuclear cataract, polar cataract, lamellar cataract, nuclear with cortical cataract, cortical cataract, sutural cataract, and total cataract (*Zhai et al., 2017*). Different types of cataracts cause different levels of visual impairment in patients. A variety of factors, including gene mutations that affect lens metabolism are closely associated with cataract (*J. Li, Chen, Yan, & Yao, 2020*), and autosomal dominant congenital cataract is the most common mode of inheritance (*Berry, Georgiou, et al., 2020*). To date, mutations in at least 50 genes involved in lens structure and development have been linked to isolated congenital cataract (*Berry, Ionides, et al., 2020; Shiels & Hejtmancik, 2017*), including crystallin genes (*CRYAA*, *CRYAB*, *CRYBB1*, *CRYBB2*, *CRYBB3*, *CRYBA1/A3*, *CRYBA2*, *CRYBA4*, *CRYGC*, *CRYGD* and *CRYGS*) (*Bhat, 2003; Zhuang et al., 2019*), membrane protein genes (*GJA3*, *GJA8*, *MIP* and *LIM2*) (*Berry, Francis, Kaushal, Moore, & Bhattacharya, 2000; Beyer, Ebihara, & Berthoud, 2013; Pei et al., 2020*), growth and transcription factor genes (*PITX3*, *MAF* and *HSF4*) (*Anand, Agrawal, Slavotinek, & Lachke, 2018*), beaded filament structural protein genes (*BFSP1* and *BFSP2*) (*Song et al., 2009*) and other genes (*CHMP4B* and *EPHA2*) (*Dave et al., 2016; Shiels et al., 2007*). Although these findings have made tremendous contributions to our understanding of the genetic etiology of congenital cataract, new disease causative genes as well as the underlying molecular mechanisms remain to be discovered (*Berry, Georgiou, et al., 2020*).

In this study, we identified a missense mutation (p.G1943E) in the phosphatidylinositol phosphate kinase (PIPK) domain of PIKfyve from a Chinese cataract family. The human *PIKFYVE* gene encodes a large protein consisting of 2,098 amino acids, which contains six evolutionarily conserved domains (*Kawasaki et al., 2012*), including zinc finger phosphoinositide kinase (FYVE) domain, plextrin homology domain, β-sheet winged helix DNA/RNA-binding motif, cytosolic chaperone CCTγ apical domain-like motif, spectrin repeats (SPEC), and C-terminal fragment PIPK domain (*S. Li et al., 2005; Shisheva, 2008*). The N-terminal FYVE domain can bind to PtdIns3P on the membrane, while the C-terminal is the kinase domain which has the catalytic function for the production of PtdIns(3,5)P_2_ (*Shisheva, Sbrissa, & Ikonomov, 1999*). The *in-vivo* function of PIKfyve in the development of lens and its association with congenital cataract have not been documented. Different from the previously reported *PIKFYVE* mutations affecting the chaperonin-like domain that cause fleck corneal dystrophy (CFD), a disease characterized by tiny white flecks scattered in corneal stroma (*Gee et al., 2015; Kawasaki et al., 2012; S. Li et al., 2005*), the p.G1943E mutation was revealed for the first time in the kinase domain of PIKfyve to cause congenital cataract in human patients. Using zebrafish models, we demonstrated that disruption of the PIPK domain in Pikfyve caused early-onset cataract defect and treating the *pikfyve*-deficient embryos with V-ATPase inhibitor bafilomycin A1 (Baf-A1) significantly alleviated the vacuolation defect in the lens of zebrafish mutants.

## Results

### Cataract family and clinical characteristics

We recruited a large Chinese Korean family affected with congenital cataract consisting of 31 family members including 12 affected individuals across 4 generations (*Figure 1A*). All the family members were Chinese Korean except one married-in individual (III-8) who was Han Chinese. There was no consanguineous marriage in this family. Bilateral cataract was observed in every generation of the pedigree and affected parents transmitted the disease to both males and females, suggesting an autosomal dominant inheritance. No other ocular and systemic abnormalities were found in these patients. Clinical information was obtained from all patients except two deceased individuals (*Table 1*). 7 of the 10 living patients were diagnosed as cataract in their first decade of life, with the smallest age of diagnosis at 2 years old. The best corrected visual acuity (BCVA) of these patients ranged from 20/125 to 20/20. The proband (III-9) showed nuclear pulverulent cataract in both eyes (*Figure 1B*) with vision loss (OD: 20/40, OS: 20/66) since childhood. The mother (II-4) of the proband also had nuclear pulverulent cataract in both eyes, while his father (II-3) had no cataract in either eye. The son (IV-5) of the proband had Y-sutural cataract. Among the other affected members of this family, individuals II-8, III-5, III-6 and IV-3 presented nuclear pulverulent opacity in both eyes, whereas individual III-1 and his son (IV-1) showed peripheral cortical punctate opacity in both eyes. All patients of this family did not show any abnormities in fundus photography and optical coherence tomography (OCT) examinations.

**Figure 1.**
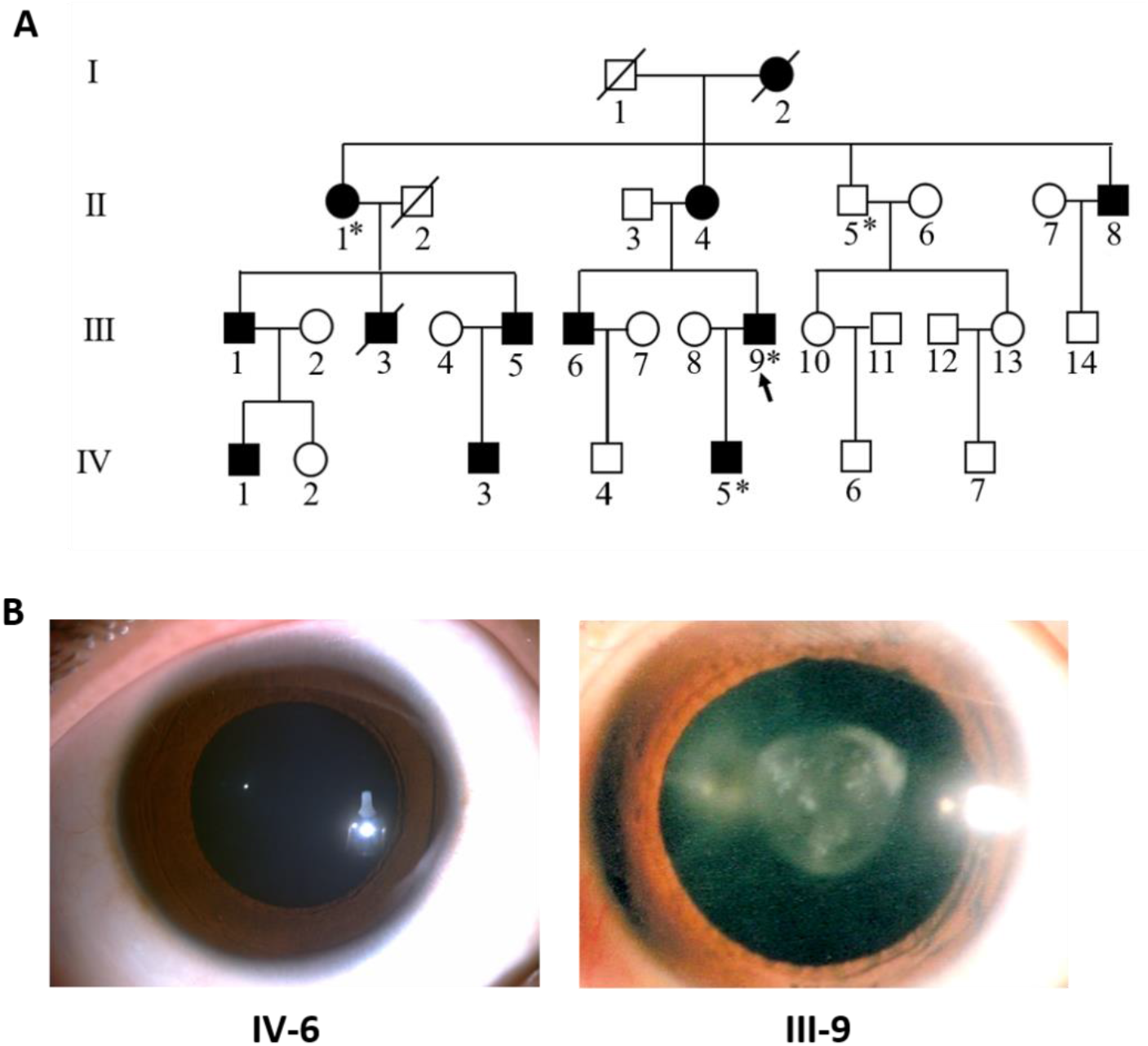
Pedigree structure and ocular manifestations of the cataract family. **(A)** Pedigree of the family with congenital cataract. Squares denote males and circles denote females; Symbols crossed by a line indicate deceased individuals. Filled symbols indicate affected individuals, while open symbols indicate unaffected individuals. All affected family members had bilateral congenital cataract. The arrow denotes the proband. The individuals marked with an asterisk (∗) are analyzed by whole exome sequencing. Genotypes of the *PIKFYVE* mutation (p.G1943E) are indicated below each symbol (+, wildtype allele; -, mutant allele). **(B)** Slit-lamp photographs showing the transparent lens of an unaffected individual (IV-6) and the nuclear pulverulent cataract in the left eye of the proband (III-9).

**Table 1.** Clinical characteristics of living patients in the cataract family.

| Patient | Gender | Age at diagnosis (years) | BCVA before surgery (OD, OS) | Cataract type (OU) | Surgery (Y/N) | BCVA after surgery (OD, OS) |
| --- | --- | --- | --- | --- | --- | --- |
| II-1 | F | 9 | — | — | Y | 20/160, 20/125 |
| II-4 | F | 12 | 20/40, 20/32 | nuclear pulverulent | N | NA |
| II-8 | M | 4 | 20/100, LP | nuclear pulverulent | Y | 20/40, FC |
| III-1 | M | 47 | 20/25, 20/20 | peripheral cortical punctate | N | NA |
| III-5 | M | 6 | 20/40, 20/40 | nuclear pulverulent | Y | 20/20, 20/20 |
| III-6 | M | 7 | 20/40, 20/50 | nuclear pulverulent | Y | 20/16, 20/16 |
| III-9 | M | 6 | 20/40, 20/66 | nuclear pulverulent | Y | 20/28, 20/25 |
| IV-1 | M | 22 | 20/20, 20/20 | peripheral cortical punctate | N | NA |
| IV-3 | M | 3 | 20/30, 20/25 | nuclear pulverulent | N | NA |
| IV-5 | M | 2 | 20/125, 20/125 | nuclear and Y-sutural | N | NA |

BCVA: best corrected visual acuity; OD: eye, oculus dexter; OS: eye, oculus sinister; OU: eye, oculus uterque; Y: yes; N: no; F: female; M: male; LP: light perception; FC: finger counting; —: unknown; NA: not applicable.

### Identification of a *PIKFYVE* missense mutation in the cataract family

To identify the causative gene mutation responsible for congenital cataract in this family, we performed whole exome sequencing (WES) for four selected family members including three affected members (II-1, III-9 and IV-5) and one unaffected member (II-5; *Figure 1A*). The average sequencing depth was 184× and 97% of the targeted regions were covered with a read depth of at least 20×. Overall, 101,852–102,401 sequence variants were identified in each individual. Nonsynonymous or splice site variants that were shared among all three affected members but were absent from the unaffected member were filtered to retain those with minor allele frequencies < 1% in the genome aggregate database (gnomAD; https://gnomad.broadinstitute.org/). We first looked at variants in known cataract genes but did not find any disease causative mutations in this family. Further filtering on in silico prediction of pathogenicity refined the candidate list to three genes (*PIKFYVE*, *NPHS1* and *FPR1*; Table 2). We then verified and evaluated these candidate variants in the whole family using Sanger sequencing. Only the p.G1943E missense mutation in the *PIKFYVE* gene completely segregated with the disease in the family (*Figure 1A*).

**Table 2.**
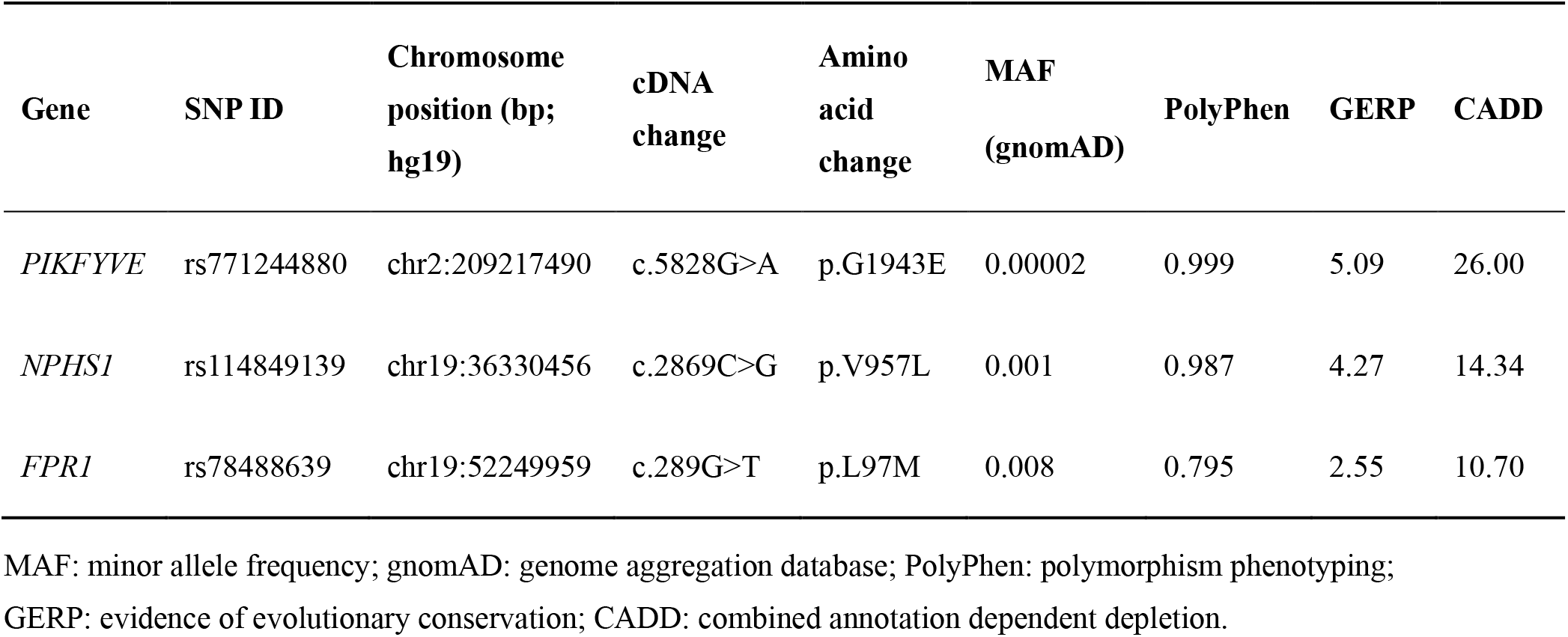
Candidate variants identified from whole exome sequencing in the cataract family.

| Gene | SNP ID | Chromosome<br>position (bp;<br>hg19) | cDNA<br>change | Amino<br>acid<br>change | MAF<br>(gnomAD) | PolyPhen | GERP | CADD |
| --- | --- | --- | --- | --- | --- | --- | --- | --- |
| *PIKFYVE* | rs771244880 | chr2:209217490 | c.5828G>A | p.G1943E | 0.00002 | 0.999 | 5.09 | 26.00 |
| *NPHS1* | rs114849139 | chr19:36330456 | c.2869C>G | p.V957L | 0.001 | 0.987 | 4.27 | 14.34 |
| *FPR1* | rs78488639 | chr19:52249959 | c.289G>T | p.L97M | 0.008 | 0.795 | 2.55 | 10.70 |

MAF: minor allele frequency; gnomAD: genome aggregation database; PolyPhen: polymorphism phenotyping; GERP: evidence of evolutionary conservation; CADD: combined annotation dependent depletion.

As shown in *Figure 2A*, in contrast to the healthy control, the *PIKFYVE* gene in the cataract patients carried a G to A heterozygous mutation in the 42^nd^ exon, which leads to the substitution of a highly conserved amino acid glycine (G) by glutamate (E) at position 1943 in the PIPK domain (*Figures 2B* and *2C*). To interpret how this mutation altered the function of PIKfyve, we first determined the protein stability of PIKfyve^WT^ and PIKfyve^G1943E^ by western blot analysis of pCS2(+)-CMV-PIKfyve^WT^ and pCS2(+)-CMV-PIKfyve^G1943E^ transfected HEK293T cells at 10 and 20 hours post transfection. As shown in *Figure 2D*, the expression level of PIKfyve^G1943E^ was comparable with that of PIKfyve^WT^, suggesting that the p.G1943E mutation has little or no effect on the protein stability of PIKfyve. Recently, a middle-to-low resolution structure of PIKfyve was resolved by cryo-EM (PDB ID:7K2V) (*Lees, Li, Kumar, Weisman, & Reinisch, 2020*). Using this structure as template, we predicted the structure of the p.G1943E mutant form of PIKfyve PIPK domain. As shown in *Figures 2E* and *2F*, the p.G1943E mutation sits in a linker connecting the N-lobe (gold) and the C-lobe (cyan) of PIPK domain. This loop (red) acts as a hinge between N-lobe and C-lobe (*Figure 2F*) and forms a highly restricted conformation. However, the p.G1943E mutation introduces a negatively charged residue to the loop. This negatively charged residue not only changes the surface electrostatic potential of PIPK domain (*Figure 2G*), but also generates charge repulsions between the loop and N-lobe (D1872 and D1871) (*Figure 2F*), thereby potentially affecting the kinase activity of PIKfyve.

**Figure 2.**
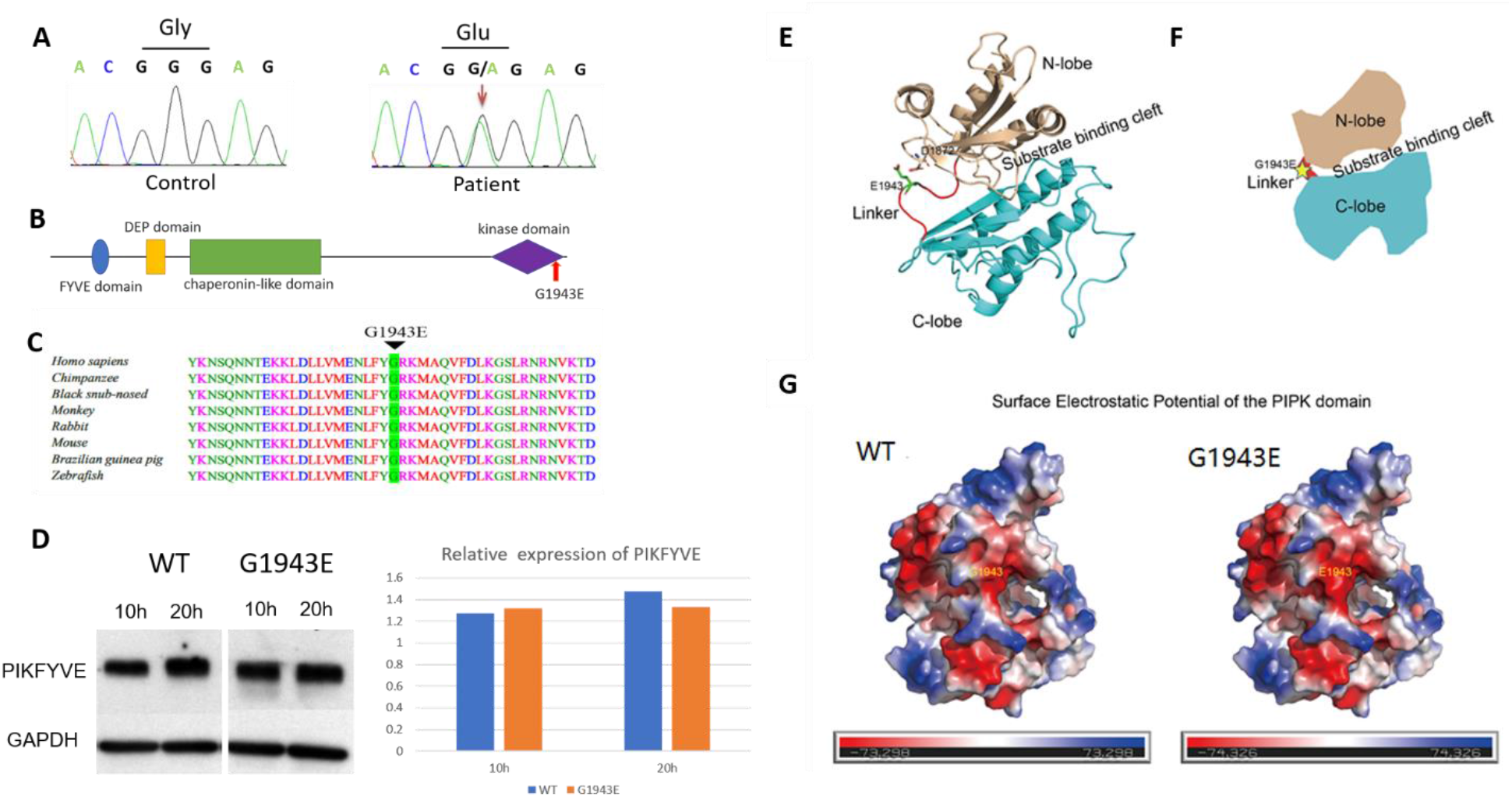
The *PIKFYVE* mutation identified from the cataract family. **(A)** Sanger sequencing chromatogram showing the cDNA sequences from a healthy control and a cataract patient. The heterozygous c.5828G>A missense mutation in the patient is indicated by red arrow. **(B)** A schematic diagram showing the human PIKfyve domains. The p.G1943E mutation in the PIPK domain is indicated by the red arrow. **(C)** Protein sequence alignment of PIKfyve orthologs in vertebrates. The black triangle denotes the conserved glycine at position 1943. **(D)** Western blot analysis of PIKfyve^WT^ and PIKfyve^G1943E^ expression in HEK293T cells that were transiently transfected with either pCS2(+)-CMV-PIKfyve^WT^ or pCS2(+)-CMV-PIKfyve^G1943E^. The protein levels were normalized by GAPDH expression. Experiments were repeated three times. **(E)** Predicted structure model of the p.G1943E mutant form of PIKfyve PIPK domain generated by the PHYPRE2 server (http://www.sbg.bio.ic.ac.uk/~phyre2/html/). N-lobe, C-lobe and the hinge linker are shown in gold, cyan and red respectively. The mutation residue E1943 is shown in sticks and labeled with green. The negatively charged residue D1872 close to E1943 side chain is also shown in sticks. **(F)** A schematic demonstrating the organization of PIKfyve PIPK domain. N-lobe, C-lobe and the hinge linker are shown in gold, cyan and red respectively. The position of the p.G1943E mutation is labeled with a yellow star. **(G)** Surface electrostatic potential comparison of the PIPK domain of PIKfyve between wild type (WT) and p.G1943E mutant. The electrostatic potentials are presented as heat maps from red to blue, and the electrostatic potential scales are shown in the lower panel. See Figure 2-source data for details.

### Disruption of PIPK domain of PIKfyve caused early-onset cataract in zebrafish

Bioinformatics analyses suggest that the p.G1943E mutation introduces a negative charged residue in the loop, which could potentially affect the kinase activity of PIKfyve. To directly demonstrate that heterozygosity of the p.G1943E mutation caused haploinsufficiency rather than dominant-negative inhibition of PIKfyve activity in human patients, we generated a *pikfyve*-deficient zebrafish mutant model using the CRISPR/Cas9-directed gene editing technology (*Shankaran, Dahlem, Bisgrove, Yost, & Tristani-Firouzi, 2017*). To maximally mimic the p.G1943E mutation in human patients, we designed single-guide RNA (sgRNA) in exon 40 to target the C terminal PIPK domain and screened out the *pikfyve^Δ8^* allele, which harbored an 8-bp deletion and a 112-bp insertion in exon 40. As shown in *Figure 3A*, this *pikfyve^Δ8^* mutation introduced a premature stop codon immediately downstream of the insertion site, thereby presumably generating a truncated Pikfyve protein that lacks the kinase activity. Phenotypical analysis revealed that *pikfyve^Δ8^* homozygous mutants developed normally before 5 days post-fertilization (dpf), but later showed severe developmental defects and all died between 7 dpf and 9 dpf (*Figure 3-figure supplement 1 A* and *B*). Under stereomicroscope, we noticed that compared to siblings (*sib*), lens of *pikfyve^Δ8^* mutants were less transparent at 5 dpf (*Figure 3B*). To characterize this phenotype in more details, we followed up the development of lens in all genotypes at different stages under differential interference contrast (DIC) microscope. Our results showed that bubble-like vacuoles first appeared in the developing lens of *pikfyve^Δ8^* homozygous mutants at 3 dpf (*Figures 3C* and *3D*). Strikingly, the number and size of vacuoles both increased drastically by 5 dpf and almost dominated the lens of mutants. Interestingly, compared to the wild-type (WT) control, *pikfyve^Δ8^* heterozygote mutants also manifested significantly higher number of vacuoles in their lens (*Figures 3C* and *3D*), suggesting that normal development of lens highly depends on the activity of Pikfyve and heterozygous disruption of Pikfyve is sufficient to cause cataract phenotype. In addition to loss of function study of Pikfyve, we also utilized a ubiquitously expressed *ubiquitin* (*ubi*) promoter to generate transgenic lines *Tg*(*ubi:PIKfyve^WT^*) and *Tg*(*ubi:PIKfyve^G1943E^*) to overexpress PIKfyve^WT^ and PIKfyve^G1943E^ in zebrafish respectively. As expected, both of these two zebrafish lines showed normal development of the lens during early stages (*Figure 3-figure supplement 1 C*). Collectively, these findings strongly supported the hypothesis that haploinsufficiency, rather than dominant-negative inhibition of PIKfyve activity by the p.G1943E mutation caused cataract in human patients. Furthermore, we noted that the vacuolation phenotype was partially rescued in *pikfyve^Δ8^;Tg*(*ubi:PIKfyve^WT^*) zebrafish (*Figure 3-figure supplement 1 D* and *E*). These findings, together with the observations that pharmacological inhibition of PIKfyve activity by inhibitor YM201636 also induced aberrant vacuolation in human lens epithelial cells HLEB3 (*Figure 3E*), suggested that function of Pikfyve in the regulation of cataract formation is highly conserved across species and our zebrafish cataract model would potentially be an excellent system for further preclinical study.

**Figure 3.**
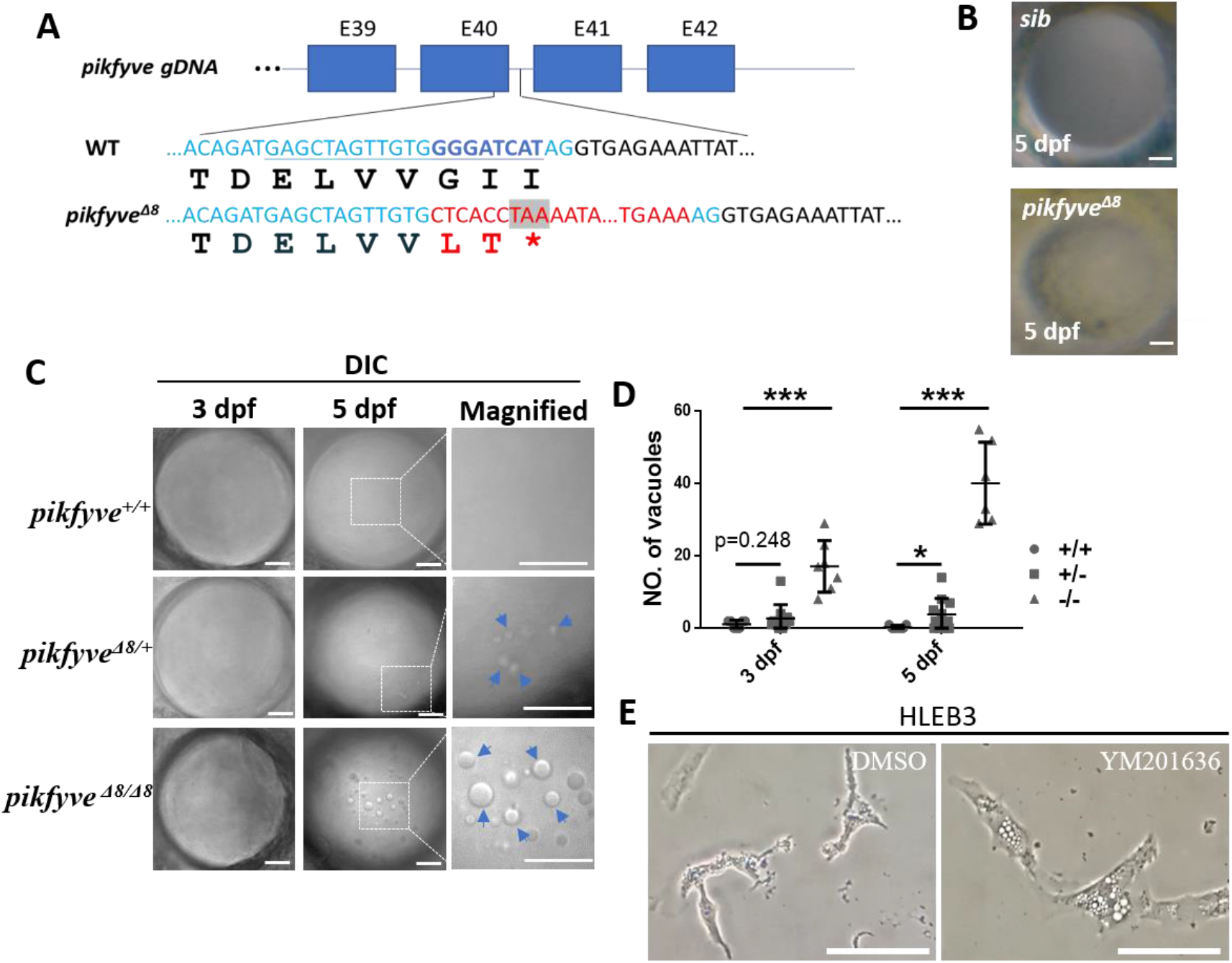
Disruption of the PIPK domain of PIKfyve in zebrafish caused early-onset cataract. **(A)** A schematic diagram showing the generated *pikfyve^Δ8^* mutant allele. The underlined base pairs are the sgRNA target. The deleted base pairs are shown in dark blue while inserted ones are shown in red. The stop codon introduced in the mutant form is shown in the grey box. **(B)** Representative images showing the lens of sibling and *pikfyve^Δ8^* mutants at 5 dpf. (**C)** Representative differential interference contrast (DIC) images showing the lens of *pikfyve*^+/+^, *pikfyve*^+/*Δ8*^ and *pikfyve^Δ8/Δ8^* embryos at 3 dpf and 5 dpf. The scale bars represent 10 μm in **(B)** and **(C)**. **(D)** Quantification of vacuole number in the lens of *pikfyve*^+/+^, *pikfyve*^+/*Δ8*^ and *pikfyve^Δ8/Δ8^* embryos at 3 dpf (n=7 for *pikfyve*^+/+^; n=10 for *pikfyve*^+/*Δ8*^; n=7 for *pikfyve^Δ8/Δ8^*) and 5 dpf (n=7 for *pikfyve*^+/+^; n=11 for *pikfyve*^+/*Δ8*^; n=6 for *pikfyve^Δ8/Δ8^*). **(E)** Representative images of HLEB3 cells treated with DMSO or PIKfyve inhibitor YM201636 for 4 hours. All experiments were repeated three times. The scale bars represent 25 μm. See Figure 3-source data for details.

### Detailed characterization of cataract phenotypes in *pikfyve^Δ8^* mutants

To delineate the details of cataract phenotype in *pikfyve^Δ8^* mutants, we crossed *pikfyve^Δ8^* mutant with the reporter transgenic line *Tg*(*cryaa:DsRed*), in which expression of DsRed was controlled by promoter of the crystalline gene *cryaa*, the major constitutive components of lens fibers. Intriguingly, in comparison with the evenly distributed DsRed signals in the lens of siblings, vacuoles in *pikfyve^Δ8^* mutants almost occupied the surface of the lens and all of them were DsRed negative (*Figure 4A*), indicating that these vacuoles did not contain lens fibers. Meanwhile, hematoxylin-eosin (HE) staining, together with ZL-1 antibody and DAPI co-staining revealed that while disruption of Pikfyve seemingly had no effects on the enucleation process of lens, lens fibers in *pikfyve^Δ8^* mutants were less organized than those in siblings (*Figures 4B* and *4C*). To further characterize the vacuoles and lens structure in high resolution, we utilized transmission electron microscope (TEM) to visualize the ultrastructure of lens in both siblings and *pikfyve^Δ8^* mutants. We could detect large vacuoles in the mutant lens at 3 dpf and 5 dpf, while only several tiny vacuoles in the sibling lens (*Figure 4D*). Lens fiber cells of WT embryos were mildly edema, and their nuclei were oval with uniform chromatin. The lens fibers (white arrows) were arranged neatly and tightly. Furthermore, mitochondria (black triangles) of WT were slightly swollen without vacuoles. In contrast, lens fiber cells of the *pikfyve^Δ8^* mutants, were obviously edematous; and the nuclei were irregularly shaped. Besides, the arrangement of lens fibers was loose and deformed. Also, lipid droplets were formed in *pikfyve^Δ8^* mutants. Compared to the WT, mitochondria (black triangles) of *pikfyve* mutants were obviously swollen and enlarged. Mitochondrial crests were reduced, and most of them were aberrantly vacuolated. In addition, autophagic lysosomes appeared swollen and vacuolated.

**Figure 4.**
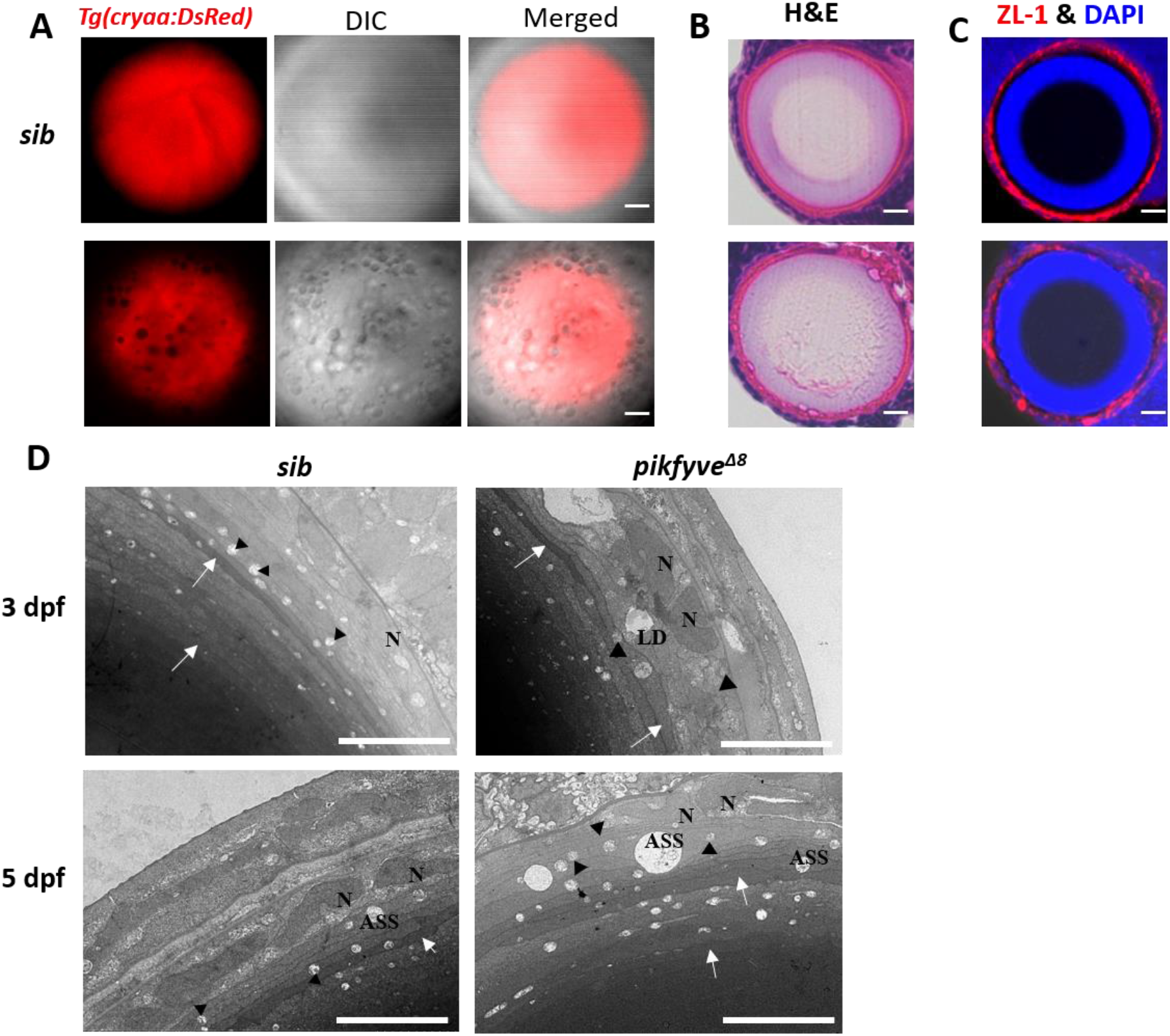
Detailed characterization of cataract phenotypes in *pikfyve^Δ8^* mutants. **(A)** Confocal imaging of the lens of 5-dpf sibling and *pikfyve^Δ8^* mutants in *Tg*(*cryaa:DsRed*) transgenic background. **(B)** Hematoxylin-eosin (HE) staining of 5-dpf sibling and *pikfyve^Δ8^* mutant zebrafish lens after cryostat section. (**C)** ZL-1 antibody staining of 5-dpf sibling and *pikfyve^Δ8^* mutant zebrafish lens. **(D)** Transmission electron microscope (TEM) images of the lens of siblings and *pikfyve^Δ8^* mutants at 3 dpf and 5 dpf. LD: lipid droplet; N: nucleus; ASS: autophagy lysosome. All results were confirmed in three different individuals. All the scale bars represent 10 μm.

### Vacuoles in *pikfyve^Δ8^* mutants were amphisomes

To define the nature of vacuoles in the lens of *pikfyve^Δ8^* mutants, we conducted time-lapse imaging to monitor their behaviors during zebrafish development. Our results showed that vacuoles in *pikfyve^Δ8^* mutants were highly dynamic and small vacuoles were frequently found to be fused with each other to form larger vacuoles (*Figure 5A,* white arrows). This feature, together with previous findings showing that PIKfyve was an essential regulator of endomembrane homeostasis (*Hasegawa, Strunk, & Weisman, 2017*), prompted us to further investigate whether vacuoles in *pikfyve^Δ8^* mutants were actually endocytic vesicles. To probe this issue, we generated fusion mRNAs encoding GFP and the markers of endocytic vesicles (i.e., the small GTPase Rab5c, Rab7 and Rab11a), which were injected into zebrafish embryos to specifically label early, late and recycling endosomes in *pikfyve^Δ8^* mutants. Our results showed that in *pikfyve^Δ8^* mutants, almost all vacuoles were positive for the late endosome maker Rab7-GFP (*Figure 5C,* white arrows). By contrast, the early endosome marker Rab5c-GFP and recycling marker Rab11a-GFP showed no co-localization with vacuoles (*Figures 5B-5D*). As autophagy was also closely-related to the organelle membrane system and had been shown to be regulated by PIKfyve (*Vicinanza et al., 2015*), we therefore checked the status of autophagosomes in *pikfyve^Δ8^* mutants by injecting *lc3b-mcherry* fusion mRNA. As shown in *Figure 5E,* a proportion of vacuoles in *pikfyve^Δ8^* mutants were also positive for the autophagosome marker Lc3b. Taken together, these data implied that vacuoles in *pikfyve^Δ8^* mutants were amphisomes formed by the fusion of autophagosome and late endosome.

**Figure 5.**
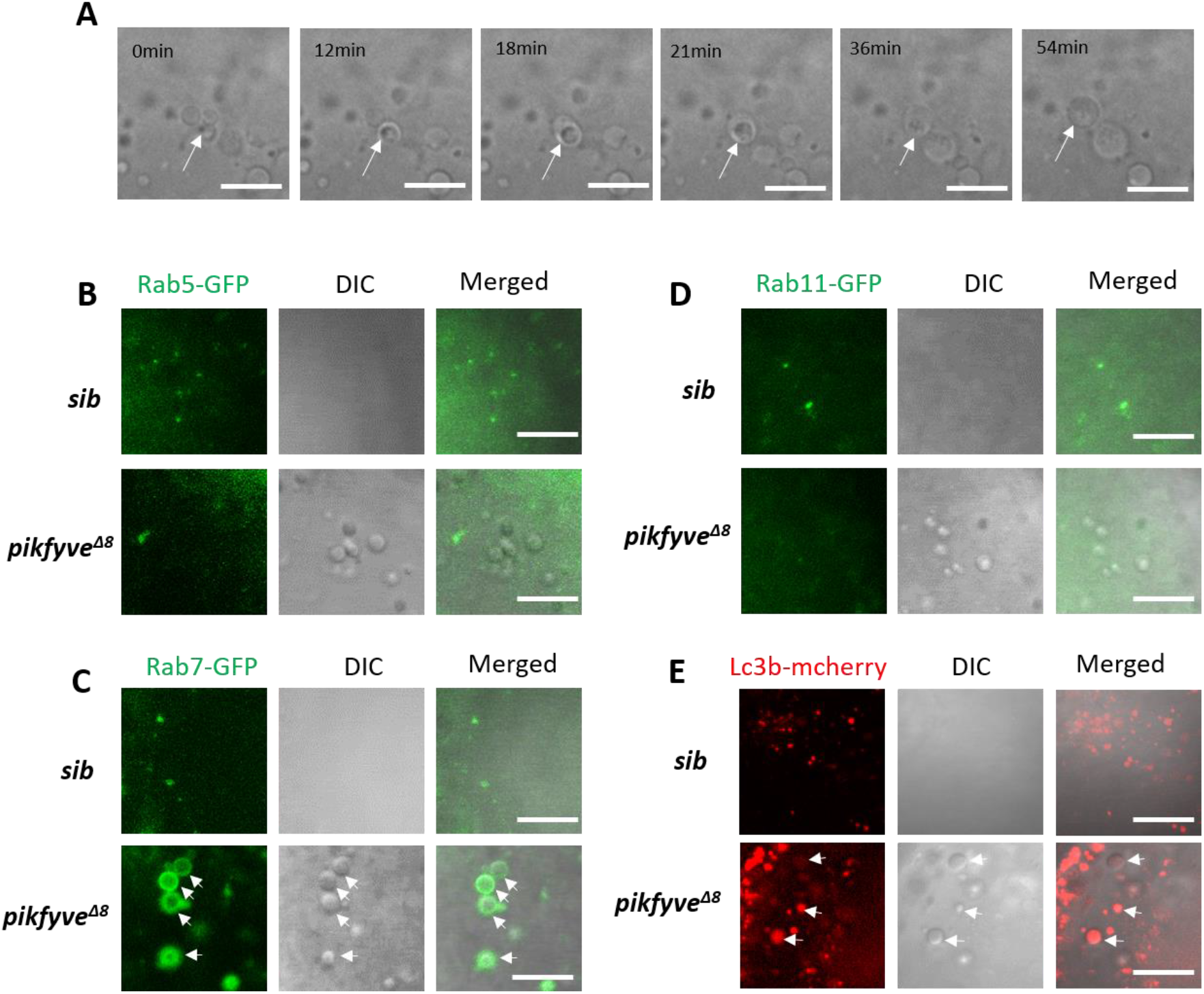
Characterization of vacuoles in *pikfyve^Δ8^* mutants. **(A)** Time-lapse imaging indicating the dynamic changes of vacuole formation in the lens of 4-dpf *pikfyve^Δ8^* mutants. White arrows indicate the fusion process of two small vacuoles. **(B)** Representative images showing the lens of 3.5-dpf sibling and *pikfyve^Δ8^* mutants injected with *gfp-rab5c* mRNA. **(C)** Representative images showing lens of 3.5-dpf sibling and *pikfyve^Δ8^* mutants injected with *gfp-rab7* mRNA. **(D)** Representative images showing lens of 3.5-dpf sibling and *pikfyve^Δ8^* mutants injected with *gfp-rab11a* mRNA. **(E)** Representative images showing lens of 3.5-dpf sibling and *pikfyve^Δ8^* mutants injected with *mcherry-lc3b* mRNA. All experiments were repeated three times. All the scale bars represent 10 μm.

### Baf-A1 partially rescued the vacuole defect in the lens of *pikfyve^Δ8^* mutant zebrafish

Baf-A1 is a specific inhibitor of V-ATPase. Previous work has shown that the vacuole phenotype induced by PIKfyve deficiency in COS-7 cells could be rescued by Baf-A1 (*Compton, Ikonomov, Sbrissa, Garg, & Shisheva, 2016*). We thus investigated whether the vacuolation defect in the lens of *pikfyve^Δ8^* mutant zebrafish could also be rescued by Baf-A1. Indeed, we found that vacuole number in the lens of *pikfyve^Δ8^* mutants treated with 1 µM Baf-A1 was significantly lower than that in the control group treated with dimethyl sulfoxide (DMSO; *Figures 6A* and *6B*). To further validate that Baf-A1 could directly alleviate cataract defect, rather than just delay the phenotype, we imaged the lens of the same mutant zebrafish before and after Baf-A1 treatment. As shown in *Figures 6C* and *6D*, while vacuole numbers of all the mutant embryos showed a slight increase after DMSO treatment or without treatment, we observed significant decrease in vacuole number in all mutants with mild or severe phenotype. On the other hand, a previous study also showed overexpression of transient receptor potential mucolipin 1 (TRPML1) could partially rescue the vacuole phenotype in *PIKfyve-deficient* macrophages (*Krishna et al., 2016*). However, in our system, ectopic expression of Trpml1 by mRNA injection failed to rescue vacuolation phenotype in *pikfyve* mutant lens (*Figure 6 – figure supplement 1*). Thus, V-ATPase rather than Trpml1 might be a key component for the formation or maintenance of the large vacuoles in the lens of *pikfyve* mutant zebrafish, and inhibition of V-ATPase by Baf-A1 could directly alleviate the cataract phenotype.

**Figure 6.**
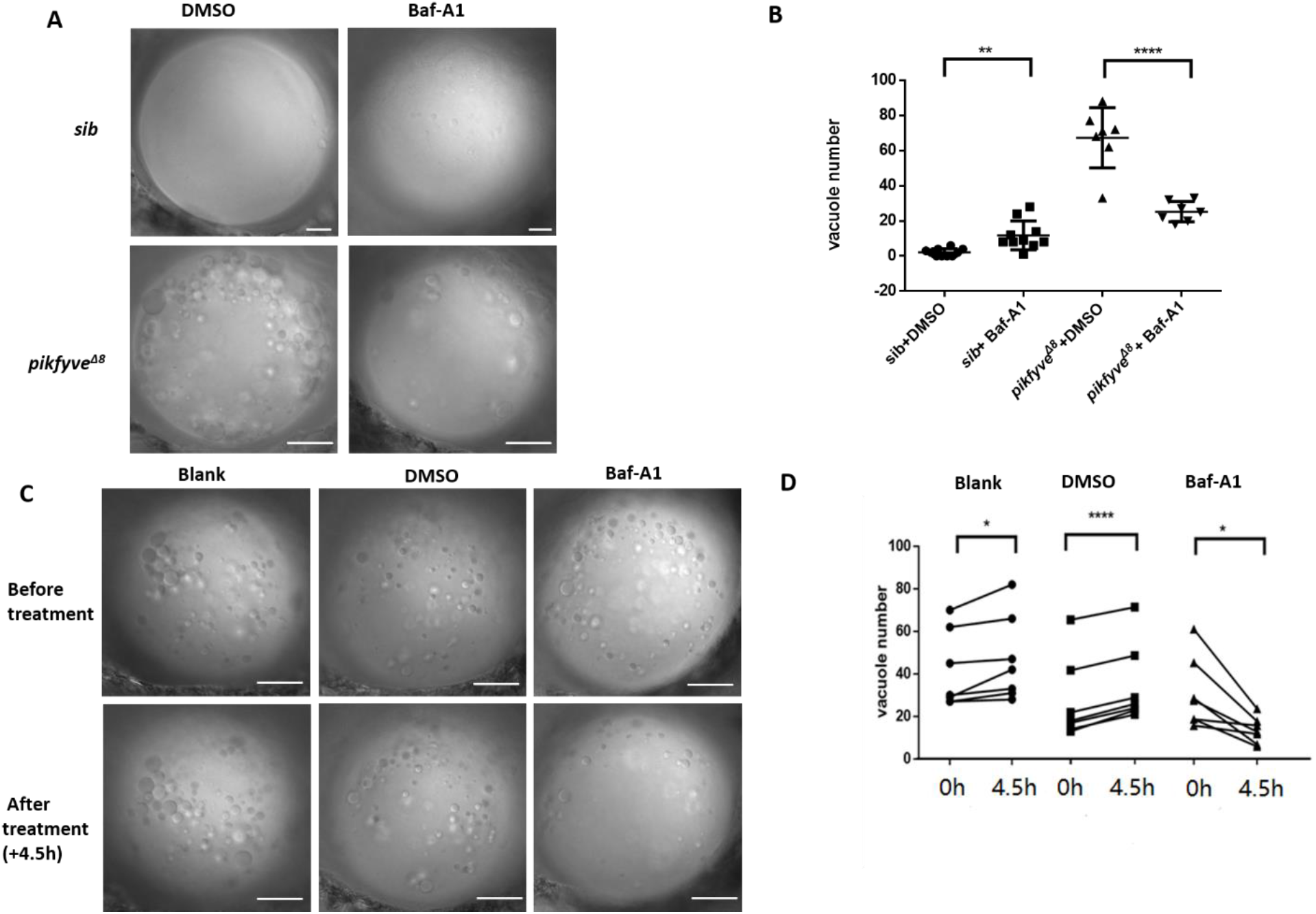
Baf-A1 partially rescued the vacuole defect in the lens of *pikfyve^Δ8^* mutant zebrafish. **(A)** Representative confocal images of the lens of 4-dpf *pikfyve^Δ8^* mutants treated with DMSO or Baf-A1 for 4.5 hours. **(B)** Quantification of the vacuole number in the lens of 4-dpf siblings and *pikfyve^Δ8^* mutant embryos treated with DMSO or Baf-A1 (n=10 for sibling groups; n=7 for mutant groups). **(C)** Confocal images of the lens of 4-dpf *pikfyve^Δ8^* mutants with no treatment or after 4.5 hours treatment with DMSO or Baf-A1. **(D)** Quantification of the vacuole number in **C** (n=6 for each group). All experiments were repeated three times. All the scale bars represent 20 μm. *, *p* < 0.05; **, *p* < 0.01; ***, *p* < 0.001; ****, *p* < 0.0001, Student’s t-test. See Figure 6-source data for details.

## Discussion

In this study, we identified a missense mutation (p.G1943E) in *PIKFYVE* responsible for congenital cataract in a Chinese Korean family (*Figures 1* and *2*). This mutation is very rare in the general population. In the gnomAD database, only 4 individuals (including 3 East Asians and 1 Latino/Admixed American) carry the p.G1943E mutation, representing an extremely low allele frequency of 0.00002 (*Table 2*). The degree and morphology of lens opacity in the cataract family were phenotypically heterogeneous (*Table 1*), which is consistent with previous findings (Berry, Ionides, et al., 2020). Most of the patients in this family developed cataract and vision loss in their childhood. Meanwhile, the majority of patients had nuclear pulverulent cataract, while the others had nuclear Y-sutural cataract or peripheral cortical punctate cataract (*Table 1*). The heterogeneity in clinical manifestations might be ascribed to interactions between genetic and environmental factors during lens development (*Berry, Ionides, et al., 2020*).

Mutations in *PIKFYVE* have been reported to be associated with CFD (*Gee et al., 2015; Kawasaki et al., 2012; Kotoulas et al., 2011; S. Li et al., 2005*). However, none of the *PIKFYVE* mutations in these studies caused congenital cataract in addition to CFD, except two of the CFD patients showed cataract formation in 66 and 58 years old respectively (*Kotoulas et al., 2011*). Interestingly, we also did not observe any corneal defect in patients with congenital cataract in this study. It is noted that all CFD related mutations in *PIKFYVE*, either homozygous or heterozygous, are distributed in two regions (i.e., amino acids 667-843 and 1490-1538), corresponding to cytosolic chaperone CCTγ apical domain-like motif and SPEC domains respectively. By contrast, the p.G1943E mutation identified in this study is located in the C-terminal PIPK domain. Based on these findings, we hypothesized that different domains of PIKfyve might exert different functions by binding with different partners. This hypothesis is further supported by other evidence. First, the predicted protein structure suggested that, instead of gross disruption of PIKfyve protein structure, the p.G1943E mutation only mildly altered the 3-D conformation of PIPK domain by changing its surface electrostatic potential and generating charge repulsions between the loop and N-lobe (*Figures 2G* and *2F*). Therefore, it is reasonable to argue that this mutation in PIPK domain may specifically alter its kinase activity or change the binding surface with other proteins, while keep the N-terminal cytosolic chaperone CCTγ apical domain-like motif and SPEC domains functional intact. Second, in our *pikfyve^Δ8^* zebrafish mutants, while the C-terminal PIPK domain was truncated in Pikfyve^Δ8^ protein, the N-terminal cytosolic chaperone CCTγ apical domain-like motif and SPEC domains were not affected. Accordingly, we did not observe any cornea defect in the zebrafish mutants either.

Inhibition of PIKfyve in COS-7 cells had been shown to induce large vacuoles through promoting the enlargement of both early and late endosomes (*Ikonomov, Sbrissa, & Shisheva, 2006; Rutherford et al., 2006*). Further studies revealed that the Ca^2+^ releasing channel, endolysosome-localized mucolipin TRPML1 acts downstream of PIKfyve to trigger membrane fusion/fission process (*Dong et al., 2010*) or promote lysosome/phagosome maturation (*Dayam, Saric, Shilliday, & Botelho, 2015; Kim, Dayam, Prashar, Terebiznik, & Botelho, 2014*). Interestingly, while overexpression of TRPML1 partially alleviated the vacuole phenotype in PIKfyve-deficient macrophages (*Krishna et al., 2016*), the rescue effect was not observed in the lens of *pikfyve^Δ8^* zebrafish mutants (*Figure 6 – figure supplement 1*), suggesting that PIKfyve might function in a context dependent manner. Consistent with this idea, we also noted that vacuoles in macrophages and lens cells of *pikfyve^Δ8^* mutants were differently stained by lysosome marker (*Figure 5 – figure supplement 1*). On the other hand, vacuole formation in PIKfyve-deficient COS-7 cells and macrophages could also be inhibited by a drug called Baf-A1 (*Compton et al., 2016; Isobe et al., 2019*). In this study, we found that Baf-A1 could partially rescue the vacuole defect in the lens of *pikfyve^Δ8^* mutant zebrafish (*Figures 6A-6D*). Baf-A1 is a macrolide antibiotic that inhibits V-ATPase, the ATP-dependent proton pump located on the membrane of organelles. The cellular acidification process mediated by V-ATPase may affect many basic biological processes, including membrane trafficking (in particular endosome maturation and fusion between autophagosomes and lysosomes (*Hammond et al., 1998; Yamamoto et al., 1998*), protein degradation and autophagy (*Bowman, Siebers, & Altendorf, 1988; Yamamoto et al., 1998; Yoshimori, Yamamoto, Moriyama, Futai, & Tashiro, 1991*). Interestingly, several works in yeast also identified mutations in V-ATPase that did not affect proton pump function, but indeed caused defects in vacuole fusion (*Strasser, Iwaszkiewicz, Michielin, & Mayer, 2011*). Moreover, another study in drosophila demonstrated that inhibition of vesicle fusion by Baf-A1 did not depend on V-ATPases, but relied on Ca^2+^ sarco/endoplasmic reticulum Ca^2+^-ATPase (SERCA) pump, the secondary target of Baf-A1 (*Mauvezin, Nagy, Juhasz, & Neufeld, 2015*). Therefore, the target of Baf-A1 and underlying mechanisms still need to be further investigated. These mechanisms will finally contribute to the potential clinical application of the drug in the treatment of congenital cataract.

## Materials and methods

### Key resources table

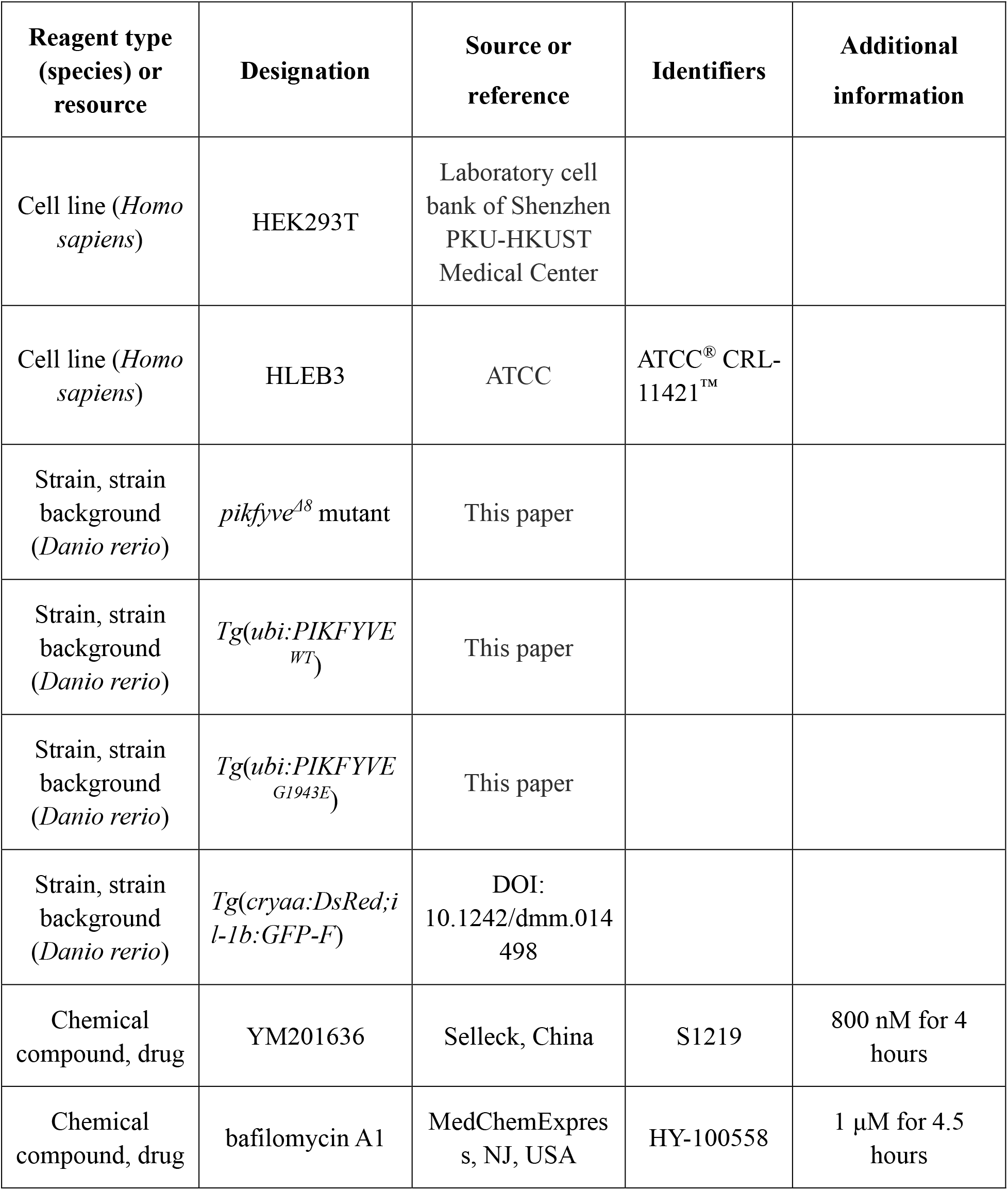
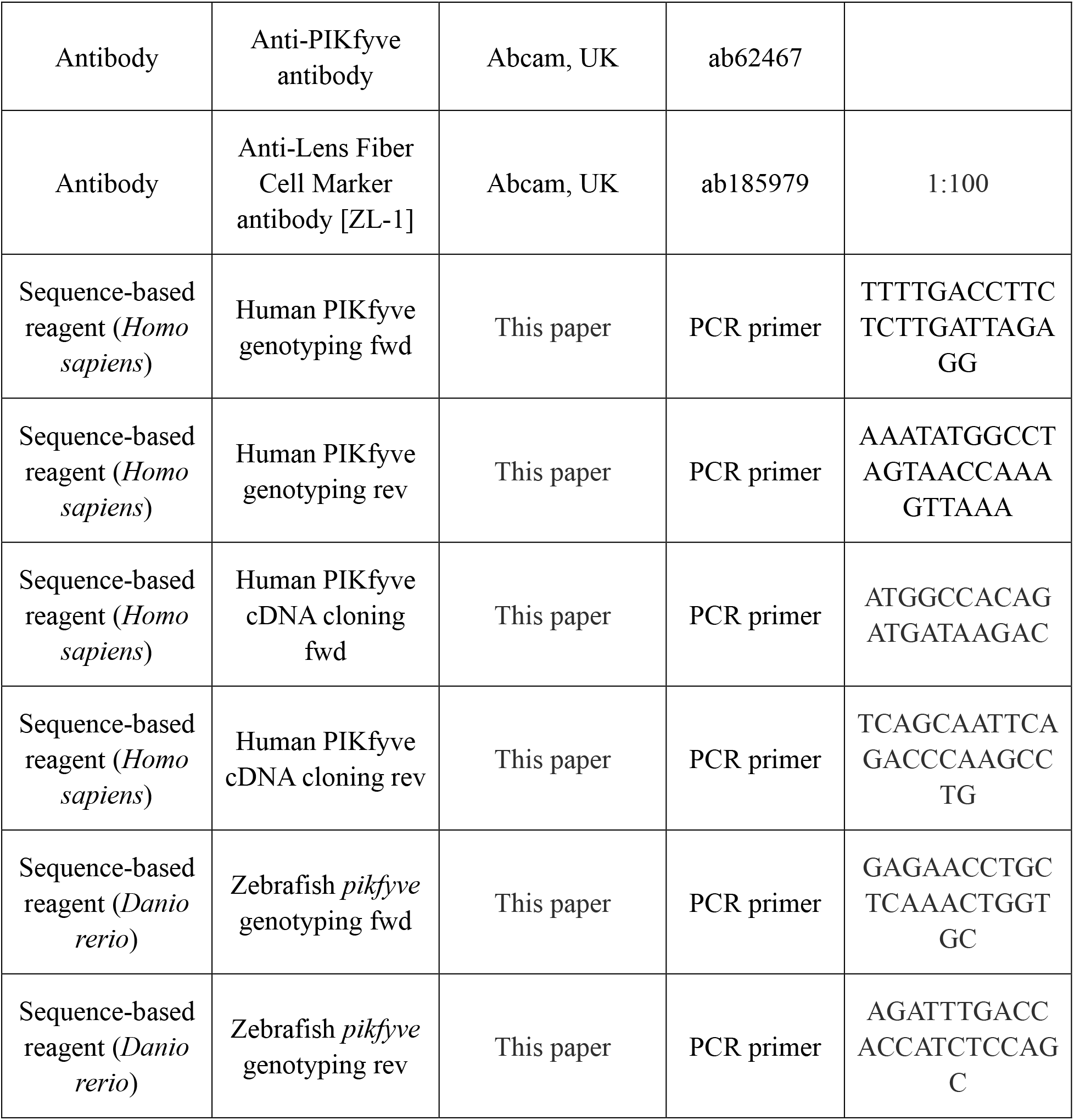

### Pedigree and patients

A 4-generation pedigree consisting of 31 family members (*Figure 1A*) was recruited from the eye clinic of Shenzhen Eye Hospital, Shenzhen, Guangdong, China. All the living family members underwent a complete ocular examination. Twelve individuals in this family were diagnosed with congenital cataract (*Figure 1B, Table 1*), while the other 19 individuals were unaffected. The diagnosis of congenital cataract was based on 1) lens opacity detected at birth or during the first decade of life; and 2) no known other cause (e.g., trauma, iatrogenic, or inflammatory disease) (Berry, Georgiou, et al., 2020). The study protocol was approved by the Independent Ethics Committee of Shenzhen Eye Hospital, in accordance with the tenets of the Declaration of Helsinki. Written informed consent was obtained from all study participants.

### Whole exome sequencing and Sanger sequencing

DNA was extracted from blood or hair samples of 21 family members (*Figure 1A*). WES was performed on 4 selected family members (II-1, II-5, III-9 and IV-5). Exome capture was performed using a SureSelect Human All Exon V6 capture kit (Agilent Technologies, Santa Clara, CA). Samples were sequenced on a HiSeq 4000 next-generation sequencing system (Illumina, San Diego, CA). Sequence reads were aligned to the human reference genome (hg19) using the Burrows-Wheeler Aligner (BWA) (*H. Li & Durbin, 2009*). Sequence variants were called using the Genome Analysis Toolkit (GATK) with the Best Practices for SNP and Indel discovery in germline DNA (*McKenna et al., 2010*). The variants were annotated using Annotate Variation (ANNOVAR) (*Wang, Li, & Hakonarson, 2010*) and then filtered based on the following criteria: 1) non-synonymous SNPs or indels in the exon region or the splice site region; 2) novel or minor allele frequency < 1% in gnomAD; 3) PolyPhen score > 0.2; 4) genomic evolutionary rate profiling (GERP) score > 2.5; and 5) combined annotation dependent depletion score (CADD) > 10. Polymerase chain reaction (PCR) and Sanger sequencing were used to verify the identified mutations from WES and screen the other family members for the mutations. The primer sequences used in PCR and Sanger sequencing were: 1) forward primer: 5’-TTTTGACCTTCTCTTGATTAGAGG-3’ and 2) reverse primer: 5’-AAATATGGCCTAGTAACCAAAGTTAAA-3’.

### Cell culture and YM201636 treatment

The HLEB3 cells (ATCC^®^ CRL-11421^™^) were obtained from ATCC and the HEK293T cells were obtained from laboratory cell bank of Shenzhen PKU-HKUST Medical Center. Cell culture was performed according to the instructions (*Andley, Rhim, Chylack, & Fleming, 1994*). The HLEB3 cells were plated into 6-well plates and grown for 4 days. The cells were then treated with YM201636 (Selleck, China) at a working concentration of 800 nM for 4 hours before imaging. Cells only treated with DMSO were used as control.

### Protein detection by western blot

Transfection of the HEK293T cells was performed according to manufacturer’s instructions of Lipofectamine RNAiMAX Reagent. Cells were transfected with 2 µg pCS2(+)-CMV-*PIKFYVE*^WT^ or pCS2(+)-CMV-*PIKFYVE*^G1943E^ and harvested after 48 hours for western blot analysis. Expression of the house keeping gene *GAPDH* was used as the loading control. The ImageJ software (National Institutes of Health, Bethesda, Maryland, USA, https://imagej.nih.gov/ij/) was used to measure the intensity of the bands and quantify the expression of PIKfyve protein. The anti-PIKfyve (ab62467, Abcam, Cambridge, UK), anti-GAPDH (ab9485, Abcam) primary antibodies and goat anti-rabbit (ab205718, Abcam) secondary antibodies were used in this study.

### Zebrafish husbandry

The AB strain zebrafish were raised and maintained according to the standard protocols (*Westerfield, 1993*). The wild type and transgenic lines including *pikfyve^Δ8^*, *Tg*(*cryaa:DsRed;il-1b:GFP-F*) (*Nguyen-Chi et al., 2014*), *Tg*(*ubi:PIKFYVE^WT^*) and *Tg*(*ubi:PIKFYVE^G1943E^*) were used in this study.

### Generation of zebrafish *pikfyve* mutant

The sgRNA was designed with the online tool Crisprscan (*Vejnar, Moreno-Mateos, Cifuentes, Bazzini, & Giraldez, 2016*). The sgRNA was synthesized by *in vitro* transcription with the MEGAshortscript™ T7 Transcription Kit (AM1354, Ambion, Austin, TX, USA) and purified by the MEGAclear™ Kit (AM1908, Ambion). The sgRNA, together with the Cas9 protein (EnGen® Cas9 NLS, S. pyogenes, M0646M, NEB, Ipswich, MA, USA) was injected into zebrafish embryos at one-cell stage. The final concentration was 100 ng/μL for each sgRNA and 800 ng/μL for Cas9 protein. Stable mutant line was screened by the T7 endonuclease 1 (T7E1) digestion and Sanger sequencing.

### Generation of transgenic zebrafish lines

The coding sequence of wild type and G1943E mutant form of human *PIKFYVE* was placed downstream of *ubi* promoter (*Mosimann et al., 2011*), which was then cloned to a modified PBSK vector containing Tol2 element (*Kawakami, Shima, & Kawakami, 2000*). The plasmid and transposon mRNA were then injected into embryos at one-cell stage. Injected embryos were raised into adulthood and screened for germline transmission by PCR and whole-mount in situ hybridization (WISH) of F1 embryos.

### Histology

Zebrafish embryos were fixed in 4% PFA at 5 dpf and then dehydrated in 30% sucrose. The whole embryos were mounted in optimal cutting temperature compound and frozen in -80°C. Cryo-section was performed at 10 µM intervals in transverse planes from the head. HE staining and antibody staining were conducted according to the standard protocols. ZL-1 antibody (ab185979, Abcam) was used in this study. TEM analysis of the lens was completed by Servicebio Technology (Wuhan, China).

### Imaging of zebrafish lens

Images were taken under ZEISS LSM980 Confocal Laser Scanning Microscope (Carl Zeiss Microscopy, NY, USA). The 63× objective (Plan-Apochromat 63×/1.40 Oil DIC M27) was used in this study. For time-lapse imaging experiment, 4-dpf zebrafish was mounted in 1.0% low-melting agarose with 0.02% tricaine. The lens was directly imaged under ZEISS Celldiscoverer 7 microscope (Carl Zeiss Microscopy). A 50× water lens (Plan-Apochromat 50x/1.2) was used and images were taken every 3 minutes. Images were analyzed by ImageJ (National Institutes of Health).

### Detection of endosomes and autophagosomes in zebrafish lens

Coding sequences of *rab5c*, *rab7* or *rab11a* were fused with *GFP* sequence and cloned into the pCS2+ vector. mRNAs were synthesized *in vitro* using the mMESSAGE mMACHINE SP6 kit (AM1908, Ambion). 1-2 nL mRNA (100 ng/μL) was injected into zebrafish embryos at one-cell stage. At 3.5 dpf, embryos were mounted in 1% low-melting agarose and imaged under the confocal microscope.

### Baf-A1 treatment

Baf-A1 powder (HY-100558, MedChemExpress, NJ, USA) was dissolved to the concentration of 1 mM in DMSO for stock, and diluted to 1 μM in egg water before use. Zebrafish embryos were treated with either 1‰ DMSO or 1 μM Baf-A1 solution at 28.5°C for 4.5 hours and washed out by egg water before confocal imaging.

### Statistical analysis

All statistical analyses were performed using the SPSS statistics software v24.0 (IBM SPSS Statistics, NY, USA). Data was represented as mean±SD. Two-tailed Student’s t-tests were used for comparisons between two groups. Statistical significance was shown as n.s., *p* > 0.05; \**p* < 0.05; \*\*\**p* < 0.001; and \*\*\*\**p* < 0.0001. Statistical plots were generated using the ggplot2 package (*Wickham, 2009*) in R (R Core Team, 2017).

## Acknowledgements

We would like to thank all family members who participated in this study. We thank Dr. Georges Lutfalla from Institut Pasteur for providing *Tg*(*cryaa:DsRed;il-1b:GFP-F*) zebrafish. We also thank Dr. Keyu Chen for the structure analysis on PIKfyve. This study was supported by Science, Technology and Innovation Commission of Shenzhen Municipality Grants (GJHZ 20180420180937076 and JCYJ20180228164400218), and Sanming Project of Medicine in Shenzhen Grant (SZSM201812090).

## Competing interests

The authors declare no competing interests.

## Figures and figure legends

**Figure 3 – figure supplement 1.**
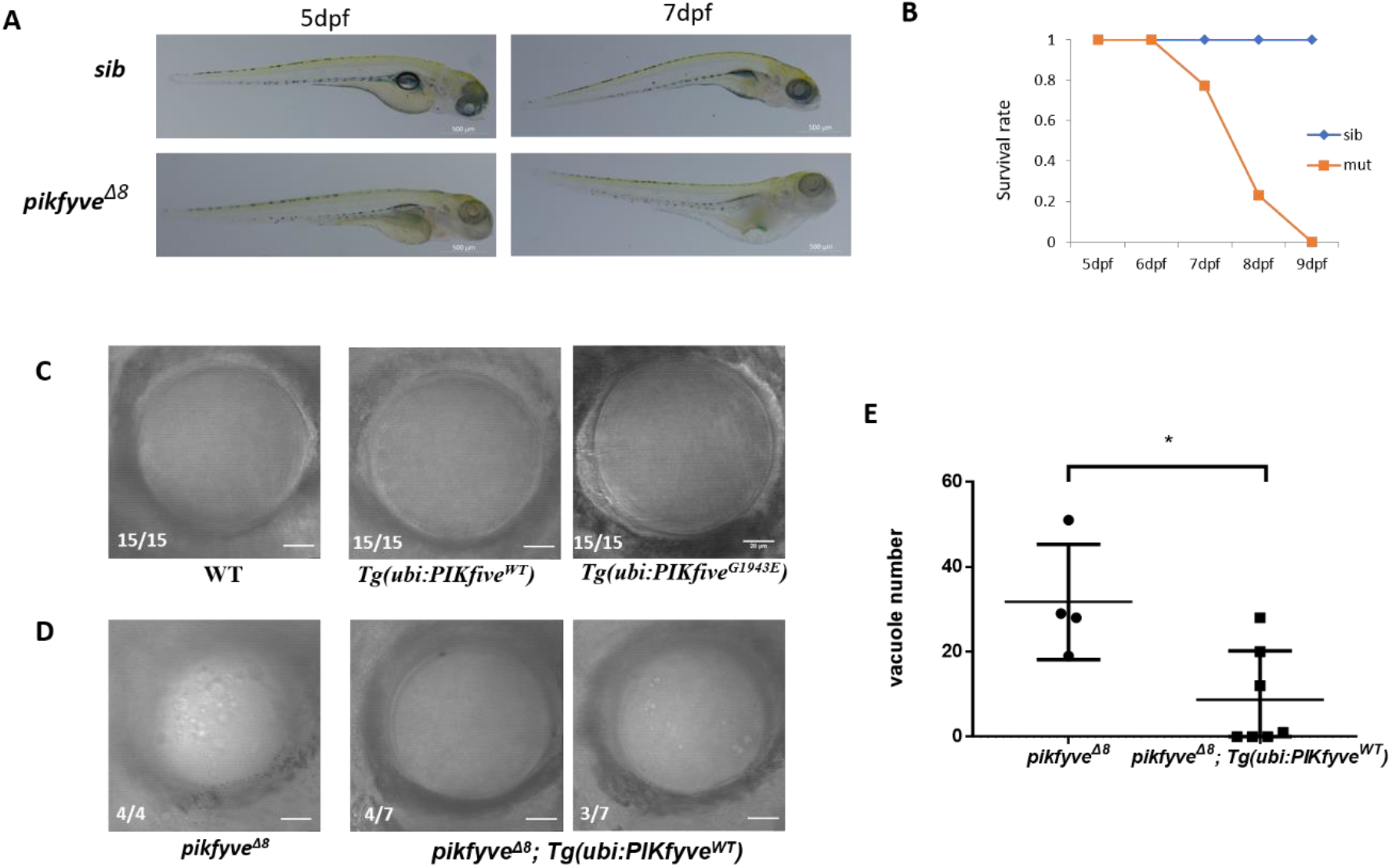
Characterization of *pikfyve-deficient* zebrafish mutants. **(A)** Gross morphology of 5-dpf and 7-dpf siblings and *pikfyve^Δ8^* mutants. The scale bars represent 500 μm. **(B)** Survival rate of *pikfyve^Δ8^* mutants (n=48) and siblings (n=50). **(C)** Representative images showing the lens of 5-dpf wild type (WT), *Tg*(*ubi:PIKfyve^WT^*) and *Tg*(*ubi:PIKfyve^G1943E^*) zebrafish embryos. (**D)** Representative images showing the lens of 5-dpf *pikfyve^Δ8^* and *pikfyve^Δ8^;Tg*(*ubi:PIKfyve^WT^*) zebrafish. The scale bars represent 20 μm in **(C)** and (**D)**. **(E)** Quantification of the vacuole number in the lens of 5-dpf *pikfyve^Δ8^* (n=4) and *pikfyve^Δ8^;Tg*(*ubi:PIKfyve^WT^*) (n=7) zebrafish embryos. *, *p* < 0.05, Student’s t-test. All experiments were repeated three times. See Figure 3 supplement 1-source data for details.

**Figure 5 – figure supplement 1.**
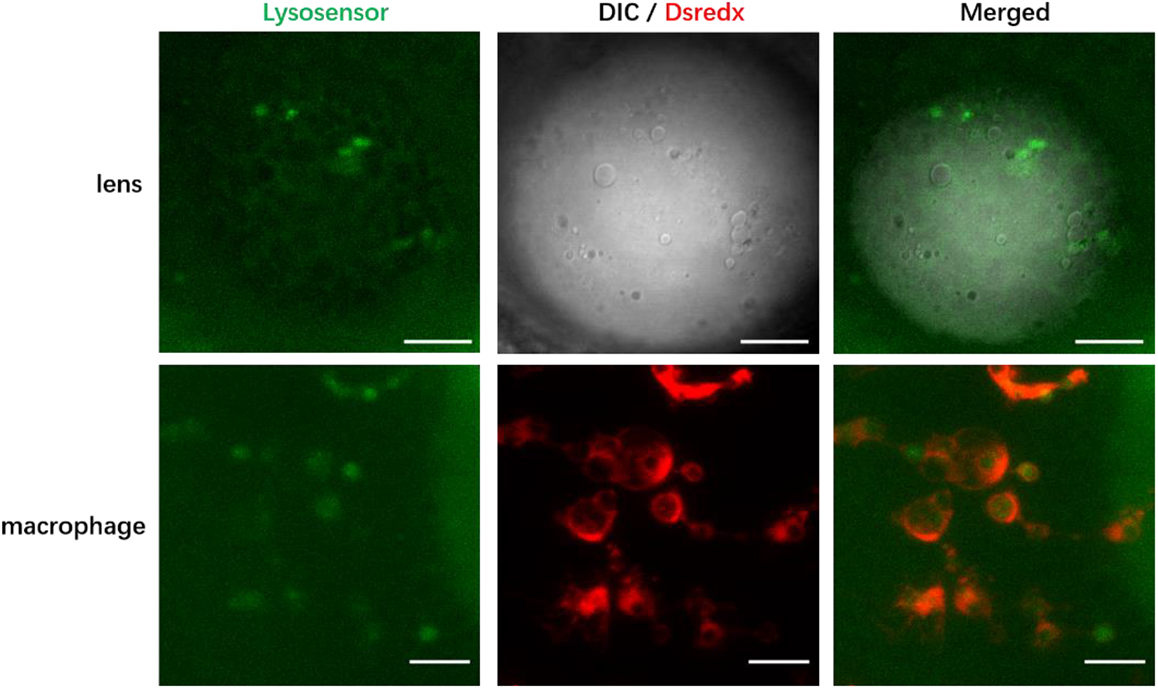
Lysosomes of *pikfyve^Δ8^* mutant zebrafish. Confocal images of the lens and macrophages of 4-dpf *pikfyve^Δ8^* mutants in *Tg*(*mpeg1:dsredx*) background after LysoSensor staining. The results were confirmed in three different individuals. The scale bars represent 20 μm.

**Figure 6 – figure supplement 1.**
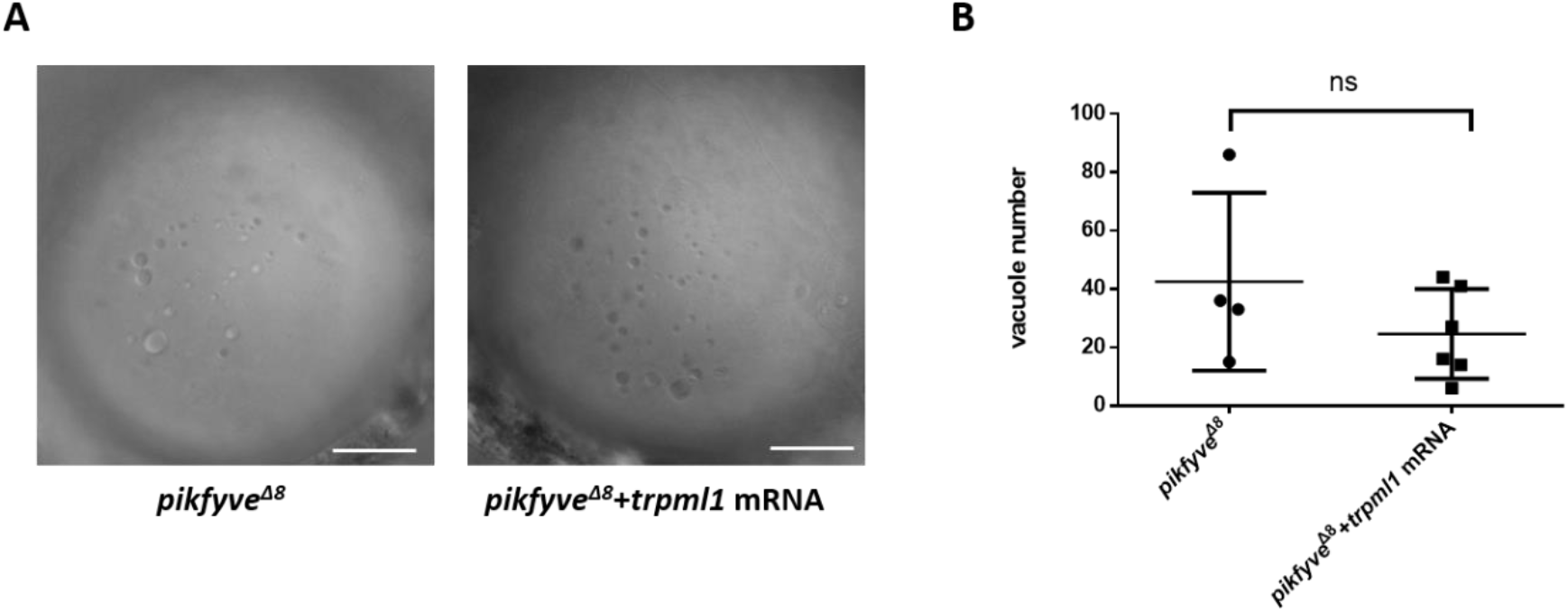
Overexpression of *trpml1* failed to rescue the lens defects in *pikfyve* mutants. **(A)** Confocal images of the lens of 4-dpf *pikfyve^Δ8^* mutants and *pikfyve^Δ8^* mutants injected with *trpml1* mRNA. The scale bars represent 20 μm. **(B)** Quantification of the vacuole number in the lens of 4-dpf *pikfyve^Δ8^* mutants (n=4) and *pikfyve^Δ8^* mutants injected with *trpml1* mRNA (n=6). n.s., not significant, *p* > 0.05, Student’s t-test. See Figure 6 supplement 1-source data for details.

## Source data

Figure 2-source data 1. Raw data for intensity of bands in Figure 2D.

Figure 2-source data 2. Raw data for the full raw unedited blots in Figure 2D.

Figure 3-source data 1. Raw data for quantification in Figure 3D.

Figure 3 supplement 1-source data 1. Raw data for quantification in Figure 3 supplement 1B and 1E.

Figure 6-source data 1. Raw data for quantification in Figure 6B and 6D.

Figure 6 supplement 1-source data 1. Raw data for quantification in Figure 6B.

## Notes

### Competing Interest Statement

The authors have declared no competing interest.

## References

Anand, D., Agrawal, S. A., Slavotinek, A., & Lachke, S. A. (2018). Mutation update of transcription factor genes FOXE3, HSF4, MAF, and PITX3 causing cataracts and other developmental ocular defects. Hum Mutat, 39(4), 471–494. doi:10.1002/humu.23395

Andley, U. P., Rhim, J. S., Chylack, L. T., Jr., & Fleming, T. P. (1994). Propagation and immortalization of human lens epithelial cells in culture. Invest Ophthalmol Vis Sci, 35(7), 3094–3102.

Berry, V., Francis, P., Kaushal, S., Moore, A., & Bhattacharya, S. (2000). Missense mutations in MIP underlie autosomal dominant ’polymorphic’ and lamellar cataracts linked to 12q. Nat Genet, 25(1), 15–17. doi:10.1038/75538

Berry, V., Georgiou, M., Fujinami, K., Quinlan, R., Moore, A., & Michaelides, M. (2020). Inherited cataracts: molecular genetics, clinical features, disease mechanisms and novel therapeutic approaches. Br J Ophthalmol, 104(10), 1331–1337. doi:10.1136/bjophthalmol-2019-315282

Berry, V., Ionides, A., Pontikos, N., Georgiou, M., Yu, J., Ocaka, L. A., . . . Michaelides, M. (2020). The genetic landscape of crystallins in congenital cataract. Orphanet J Rare Dis, 15(1), 333. doi:10.1186/s13023-020-01613-3

Beyer, E. C., Ebihara, L., & Berthoud, V. M. (2013). Connexin mutants and cataracts. Front Pharmacol, 4, 43. doi:10.3389/fphar.2013.00043

Bhat, S. P. (2003). Crystallins, genes and cataract. Prog Drug Res, 60, 205–262. doi:10.1007/978-3-0348-8012-1_7

Bowman, E. J., Siebers, A., & Altendorf, K. (1988). Bafilomycins: a class of inhibitors of membrane ATPases from microorganisms, animal cells, and plant cells. Proc Natl Acad Sci U S A, 85(21), 7972–7976. doi:10.1073/pnas.85.21.7972

Compton, L. M., Ikonomov, O. C., Sbrissa, D., Garg, P., & Shisheva, A. (2016). Active vacuolar H+ ATPase and functional cycle of Rab5 are required for the vacuolation defect triggered by PtdIns(3,5)P2 loss under PIKfyve or Vps34 deficiency. Am J Physiol Cell Physiol, 311(3), C366–C377. doi:10.1152/ajpcell.00104.2016

Dave, A., Martin, S., Kumar, R., Craig, J. E., Burdon, K. P., & Sharma, S. (2016). EPHA2 mutations contribute to congenital cataract through diverse mechanisms. Mol Vis, 22, 18–30.

Dayam, R. M., Saric, A., Shilliday, R. E., & Botelho, R. J. (2015). The Phosphoinositide-Gated Lysosomal Ca(2+) Channel, TRPML1, Is Required for Phagosome Maturation. Traffic, 16(9), 1010–1026. doi:10.1111/tra.12303

Dong, X. P., Shen, D., Wang, X., Dawson, T., Li, X., Zhang, Q., . . . Xu, H. (2010). PI(3,5)P(2) controls membrane trafficking by direct activation of mucolipin Ca(2+) release channels in the endolysosome. Nat Commun, 1, 38. doi:10.1038/ncomms1037

Gee, J. A., Frausto, R. F., Chung, D. W., Tangmonkongvoragul, C., Le, D. J., Wang, C., . . . Aldave, A. J. (2015). Identification of novel PIKFYVE gene mutations associated with Fleck corneal dystrophy. Mol Vis, 21, 1093–1100.

Hammond, T. G., Goda, F. O., Navar, G. L., Campbell, W. C., Majewski, R. R., Galvan, D. L., . . . Verroust, P. J. (1998). Membrane potential mediates H(+)-ATPase dependence of “degradative pathway” endosomal fusion. J Membr Biol, 162(2), 157–167. doi:10.1007/s002329900353

Hasegawa, J., Strunk, B. S., & Weisman, L. S. (2017). PI5P and PI(3,5)P2: Minor, but Essential Phosphoinositides. Cell Struct Funct, 42(1), 49–60. doi:10.1247/csf.17003

Ikonomov, O. C., Sbrissa, D., & Shisheva, A. (2006). Localized PtdIns 3,5-P2 synthesis to regulate early endosome dynamics and fusion. Am J Physiol Cell Physiol, 291(2), C393–C404. doi:10.1152/ajpcell.00019.2006

Isobe, Y., Nigorikawa, K., Tsurumi, G., Takemasu, S., Takasuga, S., Kofuji, S., & Hazeki, K. (2019). PIKfyve accelerates phagosome acidification through activation of TRPML1 while arrests aberrant vacuolation independent of the Ca2+ channel. J Biochem, 165(1), 75–84. doi:10.1093/jb/mvy084

Kawakami, K., Shima, A., & Kawakami, N. (2000). Identification of a functional transposase of the Tol2 element, an Ac-like element from the Japanese medaka fish, and its transposition in the zebrafish germ lineage. Proc Natl Acad Sci U S A, 97(21), 11403–11408. doi:10.1073/pnas.97.21.11403

Kawasaki, S., Yamasaki, K., Nakagawa, H., Shinomiya, K., Nakatsukasa, M., Nakai, Y., & Kinoshita, S. (2012). A novel mutation (p.Glu1389AspfsX16) of the phosphoinositide kinase, FYVE finger containing gene found in a Japanese patient with fleck corneal dystrophy. Mol Vis, 18, 2954–2960.

Khokhar, S. K., Pillay, G., Dhull, C., Agarwal, E., Mahabir, M., & Aggarwal, P. (2017). Pediatric cataract. Indian J Ophthalmol, 65(12), 1340–1349. doi:10.4103/ijo.IJO_1023_17

Kim, G. H., Dayam, R. M., Prashar, A., Terebiznik, M., & Botelho, R. J. (2014). PIKfyve inhibition interferes with phagosome and endosome maturation in macrophages. Traffic, 15(10), 1143–1163. doi:10.1111/tra.12199

Kotoulas, A., Kokotas, H., Kopsidas, K., Droutsas, K., Grigoriadou, M., Bajrami, H., . . . Petersen, M. B. (2011). A novel PIKFYVE mutation in fleck corneal dystrophy. Mol Vis, 17, 2776–2781.

Krishna, S., Palm, W., Lee, Y., Yang, W., Bandyopadhyay, U., Xu, H., . . . Overholtzer, M. (2016). PIKfyve Regulates Vacuole Maturation and Nutrient Recovery following Engulfment. Dev Cell, 38(5), 536–547. doi:10.1016/j.devcel.2016.08.001

Lees, J. A., Li, P., Kumar, N., Weisman, L. S., & Reinisch, K. M. (2020). Insights into Lysosomal PI(3,5)P2 Homeostasis from a Structural-Biochemical Analysis of the PIKfyve Lipid Kinase Complex. Mol Cell, 80(4), 736–743 e734. doi:10.1016/j.molcel.2020.10.003

Li, H., & Durbin, R. (2009). Fast and accurate short read alignment with Burrows-Wheeler transform. Bioinformatics, 25(14), 1754–1760. doi:10.1093/bioinformatics/btp324

Li, J., Chen, X., Yan, Y., & Yao, K. (2020). Molecular genetics of congenital cataracts. Exp Eye Res, 191, 107872. doi:10.1016/j.exer.2019.107872

Li, S., Tiab, L., Jiao, X., Munier, F. L., Zografos, L., Frueh, B. E., . . . Schorderet, D. F. (2005). Mutations in PIP5K3 are associated with Francois-Neetens mouchetee fleck corneal dystrophy. Am J Hum Genet, 77(1), 54–63. doi:10.1086/431346

Mauvezin, C., Nagy, P., Juhasz, G., & Neufeld, T. P. (2015). Autophagosome-lysosome fusion is independent of V-ATPase-mediated acidification. Nat Commun, 6, 7007. doi:10.1038/ncomms8007

McKenna, A., Hanna, M., Banks, E., Sivachenko, A., Cibulskis, K., Kernytsky, A., . . . DePristo, M. A. (2010). The Genome Analysis Toolkit: a MapReduce framework for analyzing next-generation DNA sequencing data. Genome Res, 20(9), 1297–1303. doi:10.1101/gr.107524.110

Mosimann, C., Kaufman, C. K., Li, P., Pugach, E. K., Tamplin, O. J., & Zon, L. I. (2011). Ubiquitous transgene expression and Cre-based recombination driven by the ubiquitin promoter in zebrafish. Development, 138(1), 169–177. doi:10.1242/dev.059345

Nguyen-Chi, M., Phan, Q. T., Gonzalez, C., Dubremetz, J. F., Levraud, J. P., & Lutfalla, G. (2014). Transient infection of the zebrafish notochord with E. coli induces chronic inflammation. Dis Model Mech, 7(7), 871–882. doi:10.1242/dmm.014498

Pei, R., Liang, P. F., Ye, W., Li, J., Ma, J. Y., & Zhou, J. (2020). A novel mutation of LIM2 causes autosomal dominant membranous cataract in a Chinese family. Int J Ophthalmol, 13(10), 1512–1520. doi:10.18240/ijo.2020.10.02

Rutherford, A. C., Traer, C., Wassmer, T., Pattni, K., Bujny, M. V., Carlton, J. G., . . . Cullen, P. J. (2006). The mammalian phosphatidylinositol 3-phosphate 5-kinase (PIKfyve) regulates endosome-to-TGN retrograde transport. J Cell Sci, 119(Pt 19), 3944–3957. doi:10.1242/jcs.03153

Shankaran, S. S., Dahlem, T. J., Bisgrove, B. W., Yost, H. J., & Tristani-Firouzi, M. (2017). CRISPR/Cas9-directed gene editing for the generation of loss-of-function mutants in high-throughput zebrafish F0 screens. Curr Protoc Mol Biol, 119, 31.39.31–31.39.22. doi:10.1002/cpmb.42

Sheeladevi, S., Lawrenson, J. G., Fielder, A. R., & Suttle, C. M. (2016). Global prevalence of childhood cataract: a systematic review. Eye (Lond), 30(9), 1160–1169. doi:10.1038/eye.2016.156

Shiels, A., Bennett, T. M., Knopf, H. L., Yamada, K., Yoshiura, K., Niikawa, N., . . . Hanson, P. I. (2007). CHMP4B, a novel gene for autosomal dominant cataracts linked to chromosome 20q. Am J Hum Genet, 81(3), 596–606. doi:10.1086/519980

Shiels, A., & Hejtmancik, J. F. (2017). Mutations and mechanisms in congenital and age-related cataracts. Exp Eye Res, 156, 95–102. doi:10.1016/j.exer.2016.06.011

Shisheva, A. (2008). PIKfyve: partners, significance, debates and paradoxes. Cell Biol Int, 32(6), 591–604. doi:10.1016/j.cellbi.2008.01.006

Shisheva, A., Sbrissa, D., & Ikonomov, O. (1999). Cloning, characterization, and expression of a novel Zn2+-binding FYVE finger-containing phosphoinositide kinase in insulin-sensitive cells. Mol Cell Biol, 19(1), 623–634. doi:10.1128/mcb.19.1.623

Song, S., Landsbury, A., Dahm, R., Liu, Y., Zhang, Q., & Quinlan, R. A. (2009). Functions of the intermediate filament cytoskeleton in the eye lens. J Clin Invest, 119(7), 1837–1848. doi:10.1172/jci38277

Strasser, B., Iwaszkiewicz, J., Michielin, O., & Mayer, A. (2011). The V-ATPase proteolipid cylinder promotes the lipid-mixing stage of SNARE-dependent fusion of yeast vacuoles. EMBO J, 30(20), 4126–4141. doi:10.1038/emboj.2011.335

Vejnar, C. E., Moreno-Mateos, M. A., Cifuentes, D., Bazzini, A. A., & Giraldez, A. J. (2016). Optimized CRISPR-Cas9 System for Genome Editing in Zebrafish. Cold Spring Harb Protoc, 2016(10). doi:10.1101/pdb.prot086850

Vicinanza, M., Korolchuk, V. I., Ashkenazi, A., Puri, C., Menzies, F. M., Clarke, J. H., & Rubinsztein, D. C. (2015). PI(5)P regulates autophagosome biogenesis. Mol Cell, 57(2), 219–234. doi:10.1016/j.molcel.2014.12.007

Wang, K., Li, M., & Hakonarson, H. (2010). ANNOVAR: functional annotation of genetic variants from high-throughput sequencing data. Nucleic Acids Res, 38(16), e164. doi:10.1093/nar/gkq603

Westerfield, M. (1993). The zebrafish book : a guide for the laboratory use of zebrafish (Brachydanio rerio). Eugene, OR: M. Westerfield.

Wickham, H. (2009). ggplot2: elegant graphics for data analysis. Springer-Verlag New York.

Yamamoto, A., Tagawa, Y., Yoshimori, T., Moriyama, Y., Masaki, R., & Tashiro, Y. (1998). Bafilomycin A1 prevents maturation of autophagic vacuoles by inhibiting fusion between autophagosomes and lysosomes in rat hepatoma cell line, H-4-II-E cells. Cell Struct Funct, 23(1), 33–42. doi:10.1247/csf.23.33

Yoshimori, T., Yamamoto, A., Moriyama, Y., Futai, M., & Tashiro, Y. (1991). Bafilomycin A1, a specific inhibitor of vacuolar-type H(+)-ATPase, inhibits acidification and protein degradation in lysosomes of cultured cells. J Biol Chem, 266(26), 17707–17712.

Zhai, Y., Li, J., Yu, W., Zhu, S., Yu, Y., Wu, M., . . . Yao, K. (2017). Targeted exome sequencing of congenital cataracts related genes: broadening the mutation spectrum and genotype-phenotype correlations in 27 Chinese Han families. Sci Rep, 7(1), 1219. doi:10.1038/s41598-017-01182-9

Zhuang, J., Cao, Z., Zhu, Y., Liu, L., Tong, Y., Chen, X., . . . Yang, J. (2019). Mutation screening of crystallin genes in Chinese families with congenital cataracts. Mol Vis, 25, 427–437.

